# Studies overestimate the extent of circadian rhythm reprogramming in response to dietary and genetic changes

**DOI:** 10.1101/2020.12.18.423465

**Authors:** Anne Pelikan, Hanspeter Herzel, Achim Kramer, Bharath Ananthasubramaniam

**Affiliations:** Institute for Theoretical Biology, Humboldt Universität zu Berlin; Institute for Theoretical Biology, Charité Universitätsmedizin Berlin; Institute for Medical Immunology, Charité Universitätsmedizin Berlin

**Keywords:** Venn diagram, high-throughput analysis, differential rhythmicity, reprogramming, remodeling

## Abstract

The circadian clock modulates key physiological processes in many organisms. This widespread role of circadian rhythms is typically characterized at the molecular level by profiling the transcriptome at multiple time points. Subsequent analysis identifies transcripts with altered rhythms between control and perturbed conditions, i.e., are differentially rhythmic (DiffR). Commonly, Venn Diagram analysis (VDA) compares lists of rhythmic transcripts to catalog transcripts with rhythms in both conditions or have gained or lost rhythms. However, unavoidable errors in the rhythmicity detection propagate to the final DiffR classification resulting in overestimated DiffR. We show using artificial experiments constructed from biological data that VDA indeed produces excessive false DiffR hits both in the presence and absence of true DiffR transcripts. We present a hypothesis testing and a model selection approaches in an R-package *compareRhythms* that instead compare circadian amplitude and phase of transcripts between the two conditions. These methods identify transcripts with ‘gain’, ‘loss’, ‘change’ or have the ‘same’ rhythms; the third category is missed by VDA. We reanalyzed three studies on the interplay between metabolism and the clock in the mouse liver that used VDA. We found not only fewer DiffR transcripts than originally reported, but VDA overlooked many relevant DiffR transcripts. Our analyses confirmed some and contradicted other conclusions in the original studies and also generated novel hypotheses. Our insights also generalize easily to studies using other -omics technologies. We trust that avoiding Venn Diagrams and using our R-package will contribute to improved reproducibility in chronobiology.

## Introduction

Circadian or near-24h rhythms are present in all kingdoms of life. These rhythms regulate critical physiological processes in many species. In eukaryotes, a gene-regulatory feedback network involving a small group of *clock genes* generates cell-autonomous circadian rhythms. Transcription factors among the clock genes subsequently drive transcript rhythms in target *clock-controlled genes* (CCGs). Insights into the widespread role of circadian rhythms at the molecular level were gained by observing the effects of genetic or environmental perturbations on clock genes and clock outputs. Transcripts, which are the proximal clock output, are easily quantified using high-throughput techniques (microarray and bulk RNA-sequencing). Therefore, almost all studies focused on the effects of altered *Zeitgebers*, such as light regime and feeding, or genotype on the circadian transcriptome.

In experiments of this kind, one or more periods of the rhythm are sampled at regular intervals under the two conditions of interest. The samples themselves might consist of pools of individuals or biological replicates. The datasets obtained are subjected to statistical analyses to identify transcripts that are *differentially rhythmic* (DiffR) between the two conditions

DiffR transcripts are commonly identified using Venn diagram analysis (VDA), as we term it: A list of rhythmic transcripts is compiled under each condition using one of many popular methods (JTKcycle (Hughes et al., 2010), RAIN (Thaben and Westermark, 2014), Harmonic Regression). The two lists are compared for overlaps and differences and the results are visualized using Venn diagrams.

This approach is inappropriate for two reasons. First, VDA seemingly finds DiffR features that are rhythmic in one condition and arrhythmic in the other. This is, of course, not all we want to know. For example, VDA overlooks transcripts that remains rhythmic but has altered circadian parameters (amplitude, phase). Second, the analysis even fails to accurately find transcripts that are rhythmic in one condition but not the other. One test of rhythmicity in each of the two conditions is necessary in the VDA, but any statistical test for rhythmicity is inherently imperfect.

Statistical tests make two kinds of errors in classifying transcripts as rhythmic or arrhythmic; false positives (arrhythmic transcript classified as rhythmic) and false negatives (rhythmic transcript classified as arrhythmic). The corresponding correct classifications are true positives and true negatives. These errors can result in non-DiffR transcripts incorrectly tagged as *hits* in the DiffR analysis and vice versa. For example, a true DiffR transcript that is a false positive in one condition and true positive in the other will be considered a DiffR miss by VDA. Similarly, a transcript that is true negative and false positive in the two datasets will be considered a DiffR hit, when it is not. (‘Hit’ and ‘miss’ refer to the algorithm’s prediction of DiffR transcripts).

Any statistical test involves trade-offs between the number of false positives and false negatives. Consequently, no choice of threshold (on the p-value or test statistic) will alleviate this misclassification. Moreover, in standard situations, false positives are stringently controlled, while false negatives are tolerated. Thus, many false negatives can be expected in both conditions. Even if some of these false negatives are true positives in the other condition, VDA results in overestimating the ‘reprogramming’. Our conclusions based on transcriptomic data also hold true for any high-throughput dataset (proteomics, metabolomics), measured under two conditions. This paper examines whether VDA overestimates the number and identity of DiffR features in circadian studies and whether the misclassification of DiffR features affects the interpretation of those studies.

We illustrate using artificial scenarios constructed from real data that the VDA does indeed perform poorly and produces too many false DiffR hits. Next, we present the two different approaches to directly compare rhythms between the two conditions and identify the four categories (and not just the three categories depicted in a Venn diagram) of pertinent DiffR transcripts. We provide the available approaches in an easy-to-use R package *compareRhythms*. We reevaluate the number and identity of DiffR transcripts in three public circadian transcriptomic datasets that used VDA and compare and contrast our interpretation with theirs. We found that the extent of ‘remodeling’ is indeed much smaller that suggested in the original studies and despite this overestimation, the VDA analyses misses many DiffR transcripts that our analysis identified. This discrepancy altered some conclusions, but confirmed others and our analysis often generated novel hypotheses overlooked previously.

## Results

### VDA overestimates the number of DiffR features

We illustrate the shortcomings of the VDA using two artificial scenarios constructed from the circadian transcriptome of the mouse liver. Mouse liver transcripts were quantified every hour for 48h (Hughes et al., 2009).

### First scenario

We divided this dataset into two datasets comprising the odd and even time points (Fig. 1A). We then compared these two datasets using VDA. Controlling the false discovery rate (FDR) at 0.05 resulted in 2296 DiffR hits or 48% of all rhythmic transcripts (Fig. 1B). By construction, this scenario contains no transcripts with altered rhythmicity and hence, *no* DiffR transcripts; the two datasets represent the same rhythms. Yet, many hits are called by VDA. With a more stringent FDR threshold for rhythmicity detection, fewer DiffR hits but also fewer rhythmic transcripts were called (Fig. 1C). In fact, more than 40% of rhythmic transcripts were incorrectly called DiffR hits across a range of thresholds (Fig. 1D).

**Figure 1:**
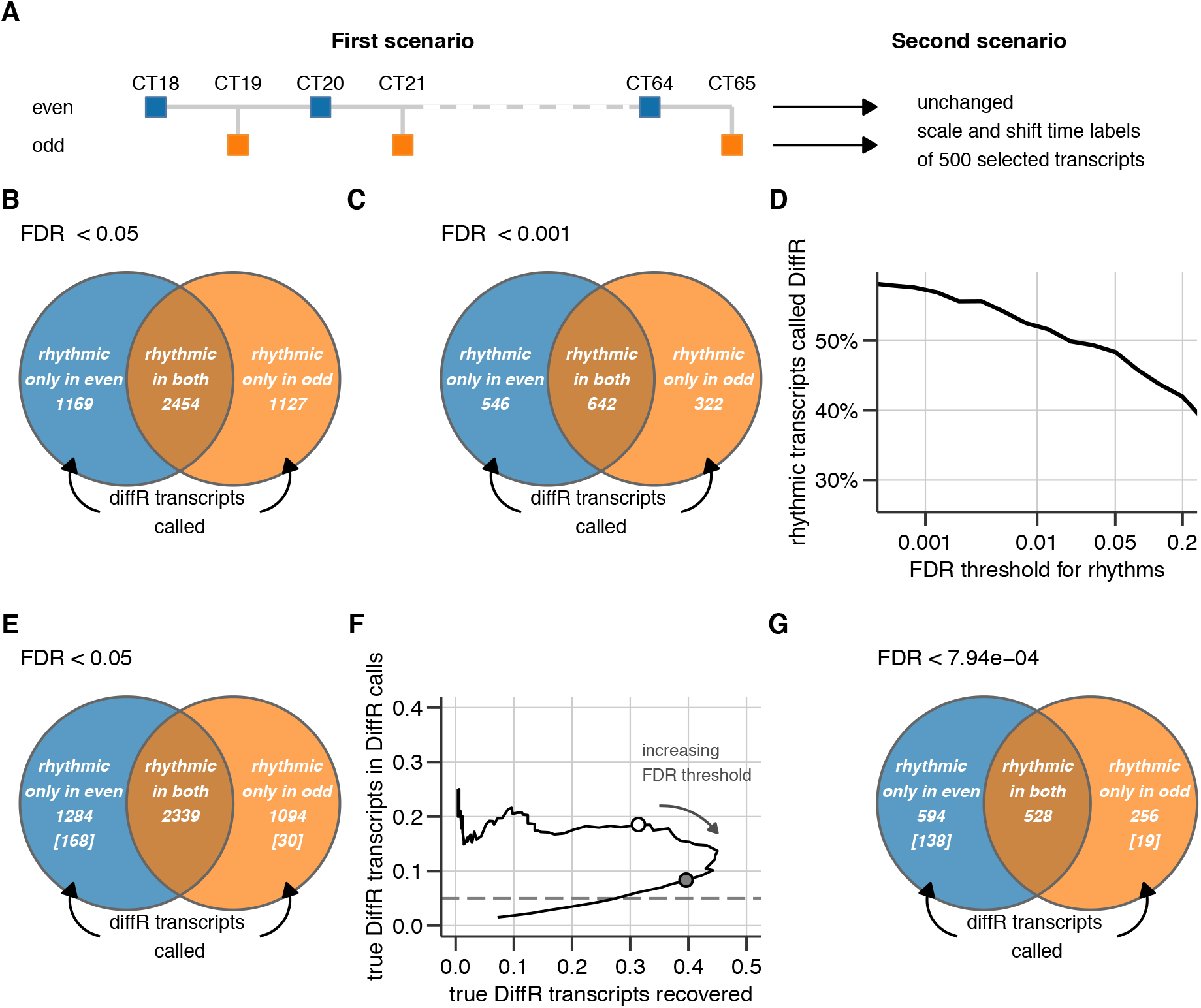
Venn diagram analysis (VDA) of two artificial circadian studies created from Hughes et al. (2009) data. (A) Construction of two scenarios from the high resolution time series of Hughes et al. (2009). Results of the VDA of the first scenario comparing old and even time samples for two different false discovery rate (FDR) thresholds: (B) 0.05 and (C) 0.001. (D) The fraction of rhythmic transcripts incorrectly identified as DiffR for different FDR thresholds under the first scenario. (E) The result of the VDA of the second scenario, where a known set of 500 transcripts is truly DiffR with changes in both amplitude and phase. (F) Precision-recall curve of the overall performance of VDA under the second scenario. The circles are two possible operating points (FDR threshold = 8 x 10^-4^(white fill), 0.05 (grey fill)). (G) VDA results for the best precision-recall performance point (open circle in (F)). The number of true DiffR transcripts identified in each group is given within square brackets.

### Second scenario

The first scenario did not include any true DiffR transcripts. Therefore, we created a second scenario with 500 true DiffR transcripts. To that end, we altered the dataset of even time points from the first scenario for these chosen 500 transcripts (Fig. 1A). We randomly altered the amplitude and phase of these transcripts and also added some noise. VDA called 2378 DiffR hits among 4717 rhythmic transcripts (or 50%) (Fig. 1E). Only a small fraction of hits (8.3%) were true DiffR transcripts. If we chose a random set of rhythmic transcripts and called them DiffR, we expect on average about 500/4717 ≈ 11% to be true DiffR transcripts in the second scenario. Thus, this random approach would outperform VDA with a standard choice of FDR threshold.

Precision-recall curves and other threshold-free measures characterize the overall performance of DiffR classification. Precision-recall curves are particularly informative when the true DiffR transcripts constitute only a small fraction of all transcripts in the data (Saito and Rehmsmeier, 2015). Precision is the fraction of true DiffR transcripts among the DiffR hits called. Recall (or Sensitivity) is the fraction of true DiffR transcripts recovered by the algorithm. Ideally, we desire methods with *both* high precision and high recall.

Precision-recall performance of VDA is poor for all choices of FDR threshold. Precision and recall of VDA was limited to 0.2 and 0.45, respectively, based on the second scenario (Fig. 1F). A precision of 0.2 and recall of 0.3 is the best combination of precision and recall (white filled circle in Fig. 1C). At this point, VDA identifies 157 true DiffR transcripts and 693 false DiffR hits (Fig. 1G). This stringent FDR threshold for rhythms (8 × 10^-4^) greatly reduces the rhythmic transcripts considered for DiffR analysis.

An amplitude threshold does not improve the performance of VDA. A minimum amplitude requirement for rhythmic transcripts helps select biologically important results (Lück and Westermark, 2016). An amplitude threshold of 0.5 log_2_ expression reduced the false DiffR hits called by VDA under the first scenario to about 40% of rhythmic transcripts (Supplemental Fig. S1A, B), but did not eliminate them for any choice of FDR threshold (Supplemental Fig. S1C). Under the second scenario, VDA with an amplitude threshold called significantly fewer DiffR hits with a reduced number of true DiffR transcripts among the hits (Supplemental Fig. S1D). The precision performance of VDA is uniform across a range of FDR thresholds (Supplemental Fig. S1E), but does not improve on the best precision-recall performance achievable. Moreover, the fewer false DiffR hits produced with an amplitude threshold comes at the cost of fewer rhythmic transcripts considered for DiffR analysis (Supplemental Fig. S1F), similar to Fig. 1G.

Thus, the presence of a large number of false DiffR hits both in data with and without true DiffR features, confirms our expectation that VDA overestimates the true number of DiffR features.

### *compareRhythms* accurately and easily finds DiffR transcripts

Clearly, a better approach than VDA is needed to accurately identify DiffR features. Two types of approaches have been proposed in the literature – hypothesis testing and model selection (Fig. 2). The hypothesis testing approach tests directly whether circadian parameters (amplitude and phase) are different between the two conditions by defining a null hypothesis for DiffR analysis (Thaben and Westermark, 2016). Rejecting this null hypothesis produces results closer to one’s intuitive understanding of DiffR features. The key steps in this approach are outlined in Fig. 2.

**Figure 2:**
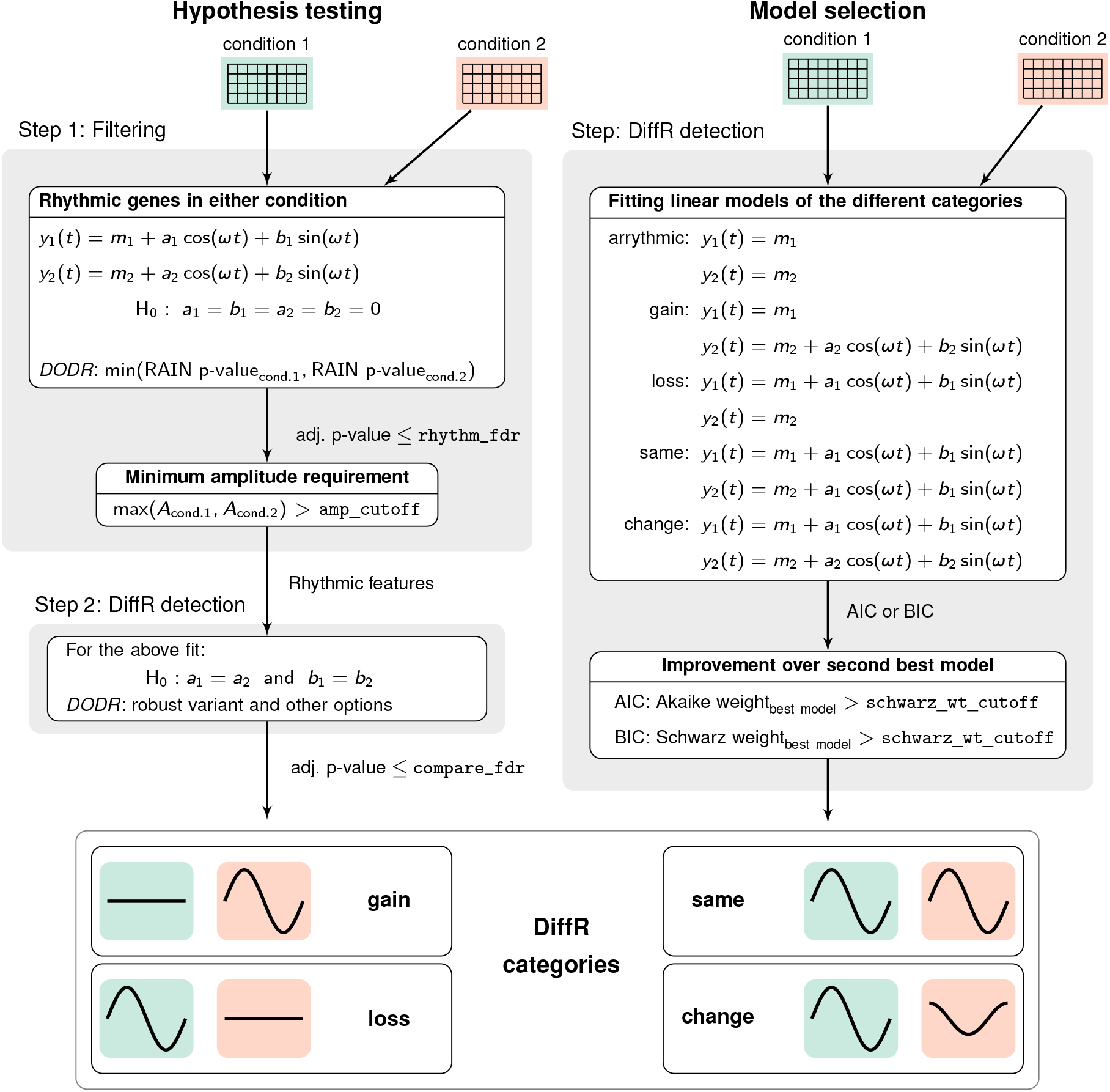
The two approaches for DiffR identification implemented in *compareRhythms*. The DiffR transcripts are classified into four categories: loss, gain, change and same rhythms in condition 2 with respect to condition 1. `rhythm_fdr, compare_fdr, amp_cutoff and schwarz_wt_cutoff` are parameters controlling different thresholds within the analysis.

Hypothesis testing produces four groups of transcripts (Fig. 2, bottom) and not just the three seen in Venn diagrams of VDA results. Let us term the two conditions A and B. Among the pre-filtered transcripts rhythmic in either A or B, there are transcripts that are (i) only rhythmic in A (and are DiffR hits), (ii) only rhythmic in B (and are DiffR hits), (iii) rhythmic in A and B (and are DiffR hits because they have different amplitude and/or phase) (iv) rhythmic in A and B (and are not DiffR hits). If A is the control condition, we could equally term these as (i) *loss* of rhythms (ii) *gain* of rhythms (iii) *change* of rhythms, and (iv) *same* rhythms. The distinction between (iii) and (iv) is nonexistent in the VDA analysis. Hence, a Venn diagram visualization is best avoided or must be altered to accommodate the fourth category.

The model selection approach (Atger et al., 2015) forgoes the choice of a null hypothesis (Fig. 2). Instead, the four rhythm groups identified by hypothesis testing along with an arrhythmic group are fit using a nested collection of harmonic regression models. The “best” model (rhythm pattern) is chosen based on an information-theoretic criterion, such as Akaike Information Criterion (AIC) or the Bayesian Information Criterion (BIC). Furthermore, the best model needs to be significantly better (threshold set by the user) than the next best model based on the same criterion.

We have implemented both these approaches in the R package *compareRhythms* (https://github.com/bharathananth/compareRhythms/). The hypothesis testing approach can be implemented using different tools for microarray data (using *limma*), RNA-seq data (using *DESeq2, edgeR* or *limma-voom*) or generic pre-normalized data (using *RAIN* and *DODR*). The model selection approach can be directly applied to any generic pre-normalized data. A single command performs the standard analysis on the two datasets by the chosen method.

Both methods in *compareRhythms* called almost no false DiffR hits in the first scenario with no true DiffR transcripts (Fig. 1A). Hypothesis testing only called the ‘same’ rhythms in 1105 transcripts (DiffR test with FDR< 0.05) and no DiffR transcripts between odd and even datasets. On the other hand, model selection called 8 *false* DiffR hits and 882 transcripts with ‘same’ rhythms (compare with Fig. 1B). Different implementations of hypothesis testing also did not find any DiffR transcripts. (By default, these methods also enforce a minimum rhythm amplitude, which is discussed further below).

Both methods recovered three-quarters of the true DiffR transcripts, but hypothesis testing had much higher precision than model selection. The analyses without an amplitude threshold aim to recover all true DiffR transcripts in the second scenario (Fig. 3, left). Hypothesis testing perfectly recalled 66% of the true DiffR transcripts (precision ≈ 1) and 75% of the true DiffR transcripts with precision above 80%. Different implementations of hypothesis testing performed identically (Supplemental Fig. S2). On the other hand, model selection suffered from a poor 50% precision in recovering 75% of the true DiffR transcripts.

**Figure 3:**
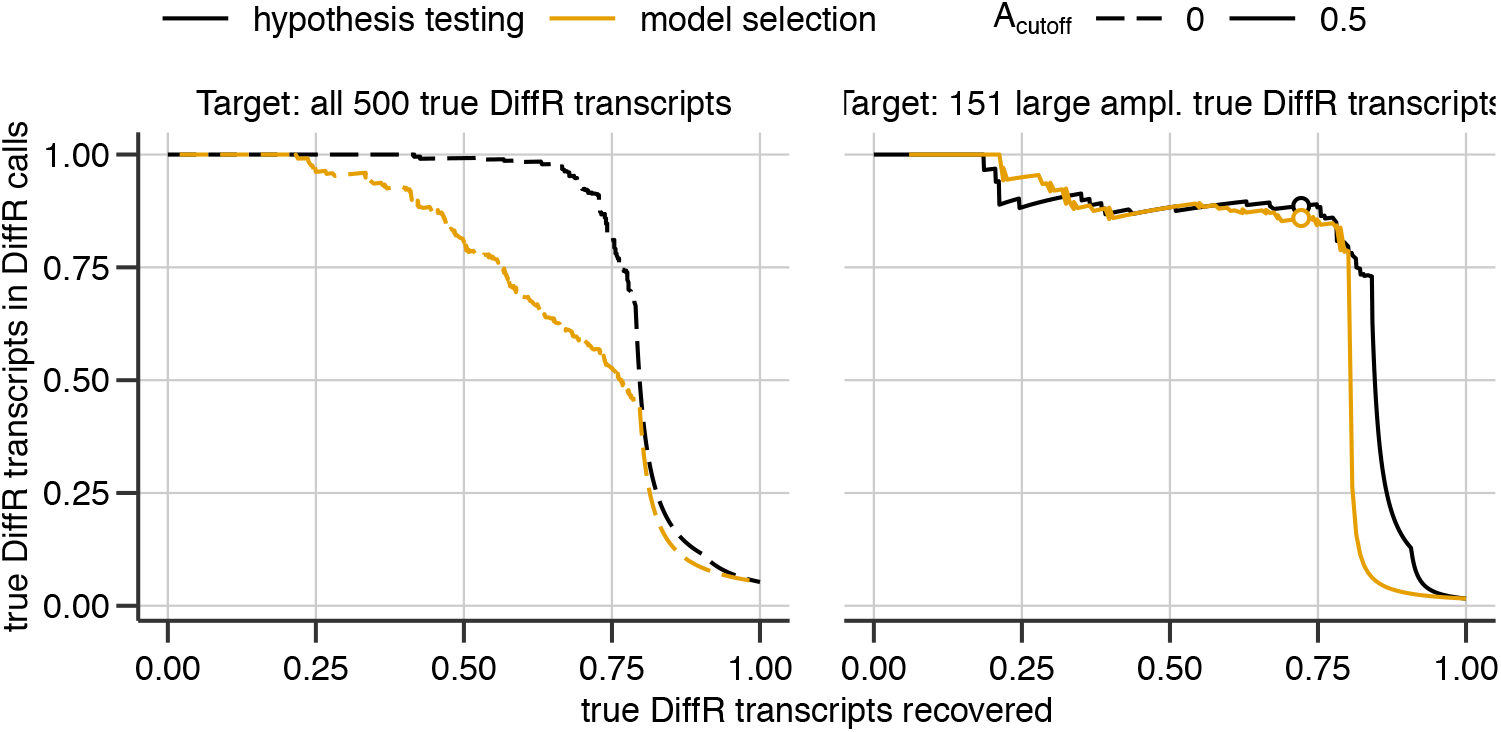
Precision-recall performance of different *compareRhythms* approaches applied to the second scenario. (*Left*) Performance of hypothesis testing and model selection on the data analyzed in Figs. 1E,F,G without an amplitude threshold (A_cutoff_) for rhythmic transcripts. (*Right*) Performance of the two approaches with an amplitude threshold on data analyzed in Fig. 1E,F,G aimed at recovering the true DiffR transcripts with biologically relevance (with amplitudes > 0.5 log_2_ expression). Performance at the default setting in *compareRhythms* is marked with circles. The curves were constructed by varying `compare_fdr` for hypothesis testing and `schwarz_wt_cutoff` for model selection.

DiffR transcripts with biological relevance based on rhythm amplitude were identified equally well by both methods. It is unclear whether all rhythmic transcripts or just those with sufficiently large amplitudes are biological meaningful. If we insist that only detection of large amplitude DiffR transcripts (with peak-to-trough amplitude > 0.5 in either group) matter, we can run both methods with an amplitude threshold (the default in *compareRhythms*). Both model selection and hypothesis testing recalled 75% of true DiffR transcripts at about 80% precision in the second scenario (Fig. 3, right), which is the performance of hypothesis testing without an amplitude threshold. Nevertheless, the amplitude threshold deteriorated the performance of hypothesis testing slightly, but improved model selection. Note, the default setting in *compareRhythms* achieves the best trade-off between precision and recall (circles in Fig. 3, right).

Although VDA clearly overestimates DiffR features, its impact on studies that used VDA is unclear. To that end, we reanalyzed three studies using *compareRhythms* to re-assess changes in rhythmicity and whether novel biological insights might have been overlooked.

### High fat diet mainly affects the core circadian clock in the liver

We re-analyzed an early high-resolution study that characterized changes in the circadian liver transcriptome in response to a nutritional challenge, i.e., high-fat diet (HFD) (Eckel-Mahan et al., 2013). We also compared these results with DiffR in response to HFD quantified using high-throughput sequencing from an independent lab (Quagliarini et al., 2019).

Hypothesis testing and model selection called less than a tenth and about a third of rhythmic transcripts, respectively, to be DiffR across both studies (Fig. 4A, Supplemental Table S1). Hypothesis testing called only 90 and 160 DiffR hits in the microarray and RNA-seq data constituting 8% and 5% of rhythmic transcripts in the respective studies. Model selection, on the other hand, called 328 and 791 DiffR hits in the two studies representing 40% and 34% of rhythmic transcripts. Model selection had lower precision (called more false DiffR hits) based on the second scenario and was expected to predict higher fractions of DiffR transcripts (Fig. 3). Interestingly, removal of the amplitude threshold reduced the fraction of DiffR transcripts, although more transcripts were called rhythmic.

**Figure 4:**
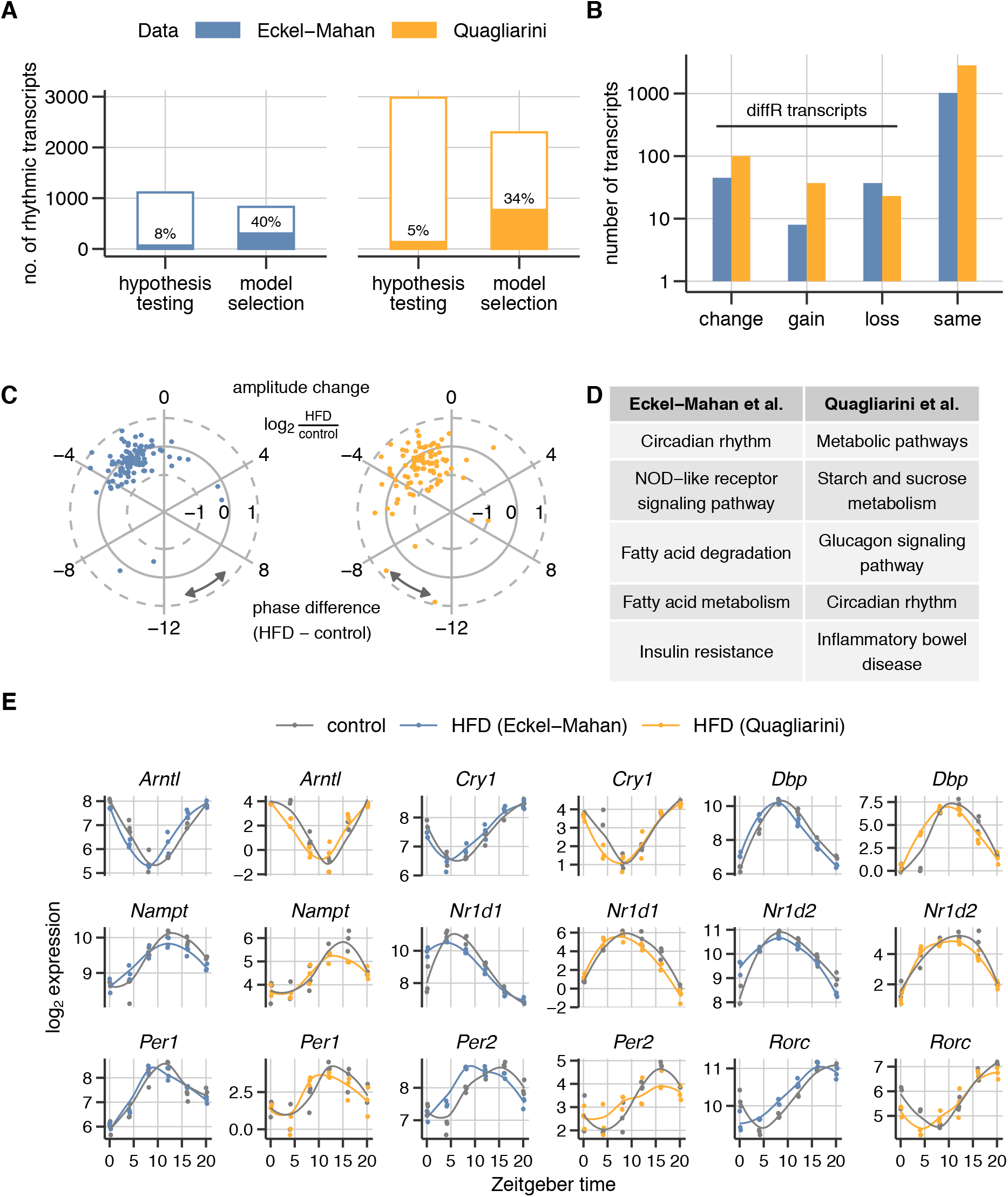
Effect of a high-fat diet (HFD) on the mouse liver clock. (A) The number of rhythmic transcripts (open bars) and DiffR transcripts (filled bars) called by the two approaches in the microarray data (Eckel-Mahan et al., 2013) and the analogous RNA-seq data (Quagliarini et al., 2019) using *compareRhythms* with default parameters. The percentage of rhythmic transcripts called DiffR is displayed within the bars. (B) The classification of DiffR hits predicted by hypothesis testing into those that ‘change’, ‘gain’ or ‘lose’ rhythms. Rhythmic DiffR misses have the ‘same’ rhythms in the two groups. (C) Circular plot representing the phase and amplitude change in the DiffR transcripts between control and HFD. Amplitude changes are represented as radial deviations from the solid gray circle and angular phase (in h) are positive for delays and negative for advances. (D) The top five KEGG enrichment categories for DiffR transcripts in each dataset. (E) The raw data as log_2_ expression for the 9 core clock genes, out of which 7 are DiffR in either the microarray or RNA-seq datasets. The lines are the mean LOESS-smoothed expression profiles for visual comparison.

DiffR hits called by hypothesis testing were a subset of those called DiffR by model selection. All but 9 transcripts called DiffR by hypothesis testing were also called DiffR by model selection in the microarray data (Supplemental Fig. S3B, Supplemental Table S1). These 9 transcripts could not be classified by model selection, since multiple models matched the expression pattern equally well. The excess DiffR hits in model selection were considered rhythmic but non-DiffR by hypothesis testing. A small number of transcripts (67) including 44 DiffR hits in model selection were considered arrhythmic by hypothesis testing. In the RNA-seq data too, DiffR hits from model selection contained the DiffR hits from hypothesis testing. Three DiffR transcripts were called arrhythmic and 35 transcripts could not be categorized by model selection due to similar performance of multiple models (Supplemental Fig. S3C, Supplemental Table S1). About 400 DiffR hits from model selection were called rhythmic but non-DiffR by hypothesis testing. This RNA-seq data contained more rhythmic transcripts according to both methods, but they did not agree on the rhythmicity of some transcripts including some that were called DiffR.

The composition of DiffR categories was similar across studies, but overlap of DiffR hits was poor. The two methods categorized DiffR transcripts similarly as ‘loss’, ‘change’ and ‘gain’ in each study (Supplemental Figs. S3B,C). The line between ‘gain’ or ‘loss’ (where transcript is not rhythmic in one condition) and ‘change’ (where transcript is rhythmic in both but with different circadian parameters) is different for the two methods and was the main source of classification differences between them. Comparing both studies, hypothesis testing predicted similar fractions of DiffR transcripts in the ‘loss’, ‘change’ and ‘gain’ categories (Fig. 4B). Differences in assays and annotations limited the number of commonly rhythmic transcripts to 736 (Supplemental Fig. S3D). Of these, only 10 transcripts were called DiffR in both, while 635 were called not DiffR in both. The remaining DiffR hits in each study were either not detected or were called arrhythmic in the other.

DiffR estimates from VDA greatly exceeded the estimates from hypothesis testing, but still missed relevant DiffR transcripts. VDA in the original microarray study (Eckel-Mahan et al., 2013) identified 2826 rhythmic transcripts, of which 2048 transcripts (72%) were called DiffR (Supplemental Fig. S3E). The RNA-seq study (Quagliarini et al., 2019), however, did not report the results of such an analysis. Only 25 of 90 DiffR hits called by hypothesis testing were also identified as DiffR by VDA in the original study (Supplemental Fig. S3F). In fact, 380 DiffR hits (19%) from VDA were predicted to have ‘same’ rhythms in both conditions and 1427 DiffR hits (70%) were considered arrhythmic by our reanalysis. As expected, VDA missed 31 transcripts that showed altered circadian parameters and classified them as rhythmic in ‘both’.

DiffR transcripts showed a consistent phase advance in HFD, however DiffR hits were not significantly enriched for any process. We observed a consistent phase advance of between 2h and 4h in almost all the DiffR transcripts across both studies (Fig. 4C). Gene enrichment analysis is often applied next to the results of the DiffR analysis to generate hypotheses. The small DiffR transcript set (Fig. 4B) was expected to make enrichment analysis less statistically powerful. Nevertheless, circadian rhythms and metabolism-related terms constituted the top five enriched KEGG categories among the DiffR transcripts (Fig. 4D); we always used all the rhythmic transcripts in either group as the background.

The individual transcript time courses were also remarkably similar between the studies (Fig. 4E). All the core clock genes in Fig. 4C except *Nr1d2* and *Per1* were DiffR in at least one study and even the two exceptions showed a trend towards earlier phases under HFD seen for the DiffR transcripts (Fig. 4C).

In summary, all the core clock gene and few CCG transcripts were DiffR under HFD with a very consistent phase advance.

### A ketogenic diet significantly activates circadian immune response in the mouse liver

We next reanalyzed microarray data on the effect of a ketogenic diet (KD) on the mouse liver (Fig. 5) and gut (Fig. S5) transcriptomes (Tognini et al., 2017).

**Figure 5:**
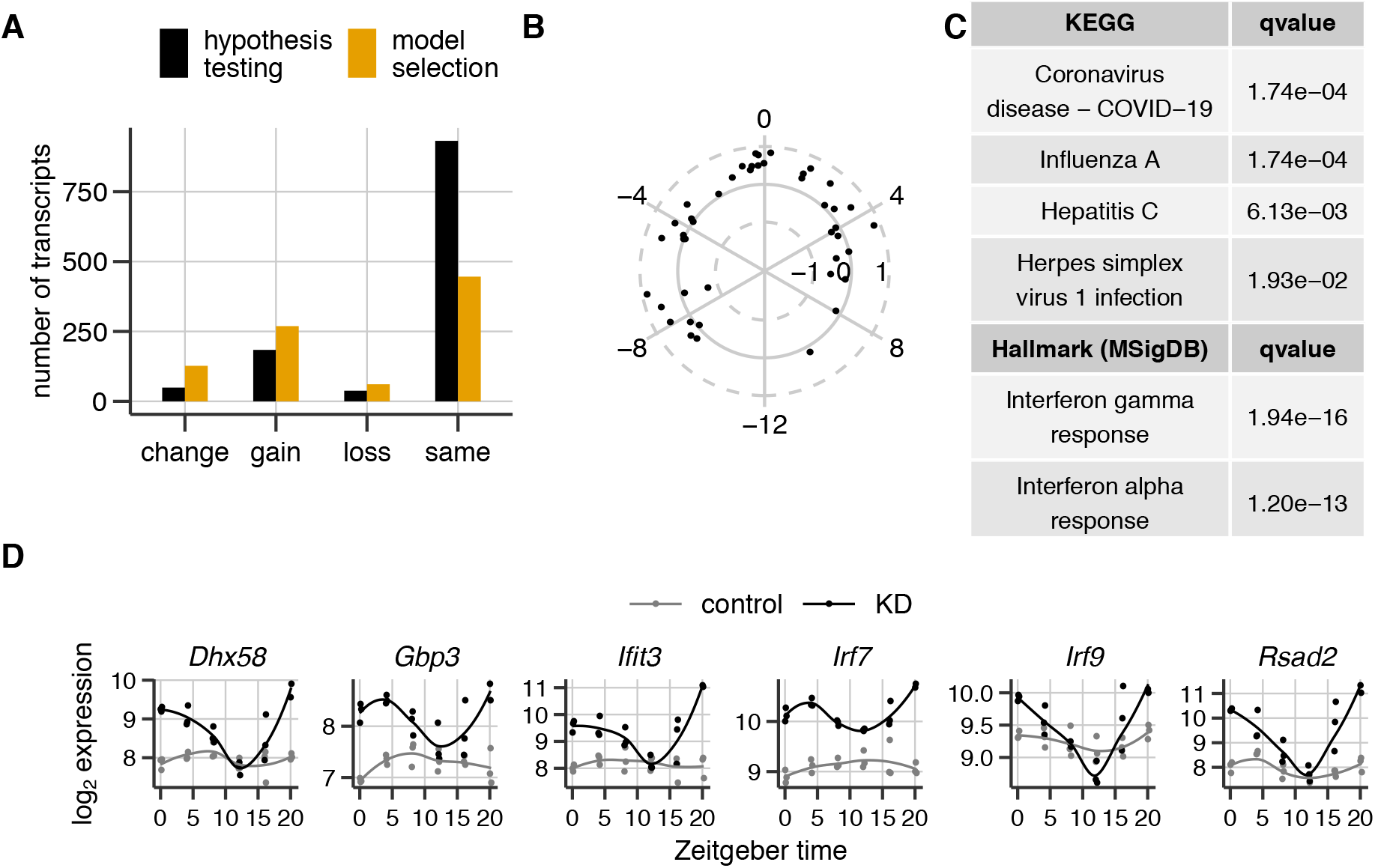
Effect of a ketogenic diet (KD) on the mouse liver clock. (A) The number of transcripts in the four categories resulting from DiffR analysis of microarray data (Tognini et al., 2017) using hypothesis testing and model selection. (B) Circular plot representing the phase and amplitude change in the transcripts in the ‘change’ category in (A) between control and KD. Amplitude changes are represented as radial deviations from the solid gray circle and angular phase (in h) are positive for delays and negative for advances. (C) KEGG and Molecular Signatures hallmark gene set enrichment of all the DiffR transcripts with the set of transcripts rhythmic in either control or KD as background. (D) Raw log_2_ expression time courses of selected transcripts involved in Interferon response under control and KD. The lines are the mean LOESS-smoothed expression profiles for visual comparison.

DiffR hits from model selection here too contained hits from hypothesis testing, and both methods predicted that most DiffR transcripts gained rhythms under KD in the liver. Hypothesis testing called 271 (23% of 1203 rhythmic transcripts) and model selection called 457 (51% of 903 rhythmic transcripts) DiffR transcripts in response to KD (Fig. 5A, Supplemental Table S2). The fraction of rhythmic transcripts called DiffR decreased without an amplitude threshold, despite increases in number of hits and number of rhythmic transcripts (Supplemental Fig. S4A). Except for 32 transcripts that could not reliably categorized by model selection (since multiple models matched data well), all DiffR hits from hypothesis testing were called DiffR by model selection. Moreover, the common DiffR hits were categorized identically. Of the additional DiffR transcripts called by model selection, 62 were called arrhythmic by hypothesis testing and 156 were considered to have the ‘same’ rhythms in the two conditions. Nevertheless, the proportions of different categories of DiffR were similar between the two methods.

VDA again missed many DiffR hits called by hypothesis testing, despite estimating large numbers of DiffR transcripts. VDA predicted 79% of rhythmic transcripts (3058 out of 3859) to be DiffR (Supplemental Fig. S4B). 75% of the DiffR transcripts called by VDA also showed novel rhythms under KD (as in Fig. 5A). Nonetheless, only 151 of the 271 DiffR transcripts called by hypothesis testing were among the hits called by VDA. Hypothesis testing placed about 40% of DiffR hits missed by VDA (as rhythmic in ‘both’) in the ‘change’ category, a category absent in the VDA.

DiffR transcripts generally increased amplitude under KD and were enriched for immune response pathways. Along with the majority of DiffR transcripts placed in the ‘gain’ category, DiffR transcripts in the ‘change’ category also showed an rhythm amplitude increase but no clear trend in the phase change (Fig. 5B). Rhythm parameter changes cannot be estimated accurately when the transcript is arrhythmic in one condition (‘gain’ and ‘loss’ categories). The significantly enriched KEGG pathways were all associated with responses to infectious disease - Influenza A, Hepatitis C and Herpes (Fig. 5C). In agreement, the DiffR transcripts were highly over-represented for hallmark genes up-regulated in response to Interferon *α* and *γ* according to the MSigDB database (Liberzon et al., 2015). Moreover, most immune response associated DiffR transcripts acquired rhythmicity under KD (Fig. 5D and Supplemental Table S2).

Fewer DiffR transcripts were present in the gut (intestinal epithelial cells) compared to the liver. Hypothesis testing and model selection called 98 (15% of 675 rhythmic transcripts) and 240 (43% of 559 rhythmic transcripts), respectively (Supplemental Fig. S5A, Supplemental Table S2). A smaller fraction of rhythmic transcripts were DiffR in the gut. As with the liver, VDA called 78% of the transcripts DiffR in the original study. Finally, the DiffR transcripts were equally distributed among the ‘gain’, ‘loss’ and ‘change’ categories of DiffR transcripts.

DiffR transcripts in ‘change’ category were phase delayed and no KEGG pathway was enriched for DiffR. ‘Change’ category transcripts were phase delayed by about 2-3 h under KD (Supplemental Fig. S5B). ‘Terpenoid biosynthesis’ was the only significant KEGG pathway enriched in the DiffR transcripts, but circadian rhythms and metabolism were present in the top three (Supplemental Fig. S5C). Nevertheless, ‘Cholesterol homeostasis’ hallmark genes were enriched among the DiffR transcripts. Many core clock genes were DiffR (Supplemental Fig. S5D), but nuclear receptors *Pparα* and *Pparγ* had indistinguishable rhythms in control and KD.

In a nutshell, KD induced more rhythm changes in the liver than HFD including *de novo* rhythms in a large subset of DiffR transcripts. These DiffR transcripts were closely associated with immune response pathways. On the other hand, rhythm changes in the gut in response to KD were limited.

### Disruption of endogenous H_2_O_2_ rhythms activates circadian oncogenic signaling

We reanalyzed finally RNA-seq data (Pei et al., 2019) on the effect of disrupting endogenous H_2_O_2_ rhythms by knocking-out (KO) *p66^Shc^* on the circadian liver transcriptome.

Both methods similarly categorized DiffR transcripts with hypothesis testing hits included in model selection hits yet again. Hypothesis testing and model selection called 424 (9% of 4483 rhythmic transcripts) and 1522 (41% of 3742 rhythmic transcripts) DiffR transcripts (Fig. 6A, Supplemental Table S3). This result was insensitive to the implementation of hypothesis testing (not shown). Here too, removal of the rhythm amplitude requirement reduced the fraction of rhythmic transcripts that were DiffR (Supplemental Fig. S6A). As before, all but 88 DiffR transcript calls by hypothesis testing agreed with model selection; 71 of these could not be reliably classified by model selection (Supplemental Fig. S6B). About 70% of the excess DiffR hits from model selection were classified as having ‘same’ rhythms by hypothesis testing. Nevertheless, the proportion of the different categories was conserved across approaches, with most DiffR hits in ‘change’ followed by ‘gain’ and then ‘loss’.

**Figure 6:**
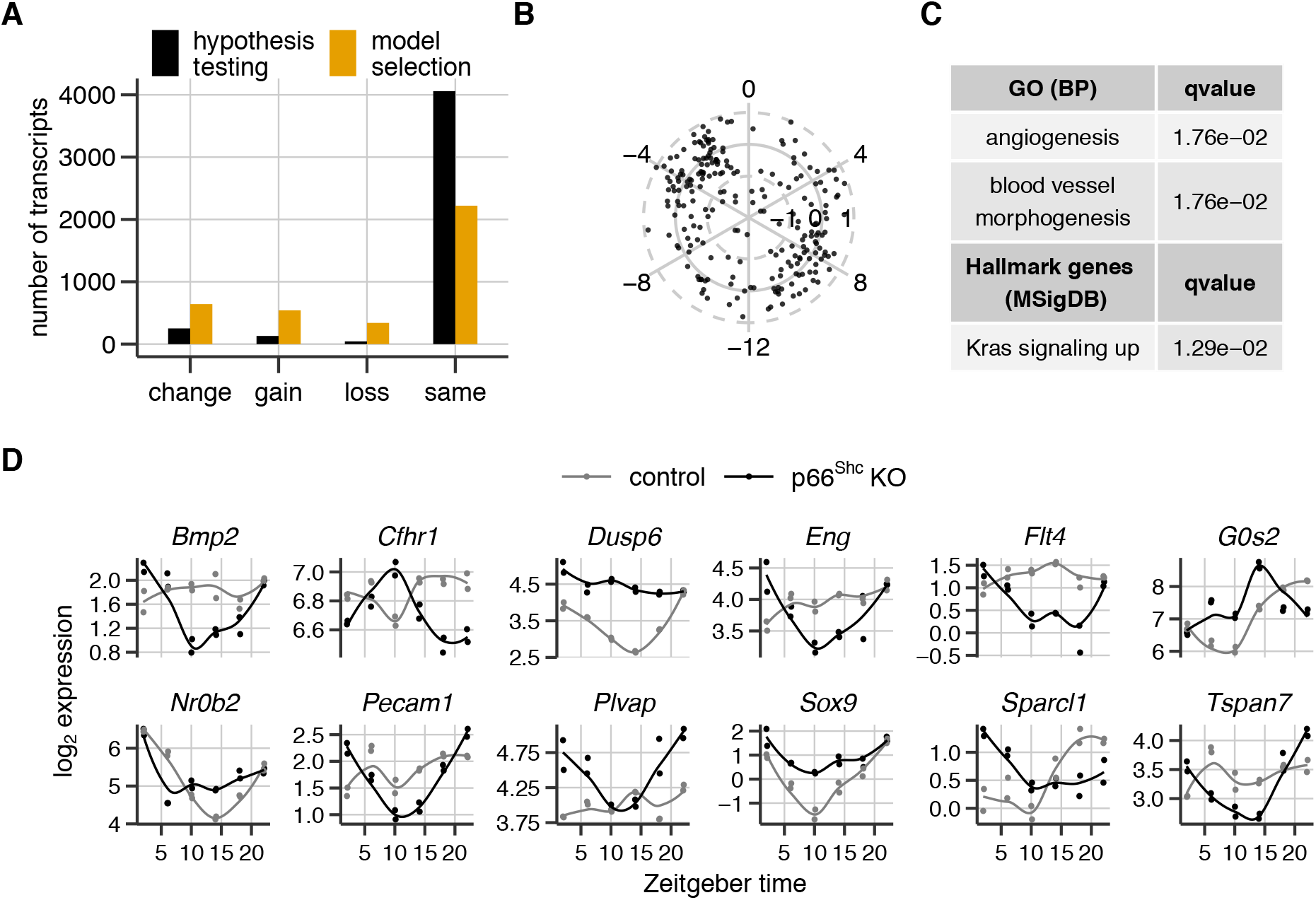
Effect of *p66^shc^* knockout (KO) on the mouse liver clock. (A) The number of transcripts in the four categories resulting from DiffR analysis of the data (Pei et al., 2019) using hypothesis testing and model selection. (B) Circular plot representing the phase and amplitude change between control and KO in the transcripts in the ‘change’ category in (A). Amplitude changes are represented as radial deviations from the solid gray circle and angular phase (in h) are positive for delays and negative for advances. (C) KEGG and Molecular Signatures hallmark gene set enrichment of all the DiffR transcripts with the set of transcripts rhythmic in either control or *p66^Shc^* KO as background. (D) Raw log_2_ expression time courses under control and *p66^Shc^* KO of selected transcripts up-regulated in KRAS signaling. The lines are the mean LOESS-smoothed expression profiles for visual comparison.

We observed a large mismatch between VDA in the original study and hypothesis testing. According to VDA in the original study, 83% of the rhythmic transcriptome (2082 of 2502 rhythmic transcripts) was DiffR (Supplemental Fig. S6C). Two thirds of the hypothesis testing DiffR hits were missed (Fig. S6D). Of these 230 transcripts were not expressed or were called arrhythmic by the VDA. Surprisingly, more than 2500 transcripts with the same rhythms in both conditions were not expressed/called rhythmic in the VDA analysis.

DiffR transcripts were enriched in vasculature development. The phase and amplitude shifts in the ‘change’ DiffR transcripts did not show a specific tendency (Fig. 6B). However, there appeared to be two cluster of phase shifts: those that are phase advanced by ~4h and those that are phase delayed by ~8h. The DiffR set was significantly over-represented for the GO categories ‘angiogenesis’ and ‘blood vessel morphogenesis’ (Fig. 6C). ‘Genes upregulated by KRAS signaling’ were also significantly enriched in the DiffR set. Most genes that overlapped with this hallmark set gained rhythms in the knockout (Fig. 6D).

To sum up, altering the endogenous H_2_O_2_ rhythms cause gain, loss and change in rhythms in the liver and these DiffR transcripts are associated with the circulatory system development and oncogenic KRAS signaling

## Discussion

High-throughput time-series profiling under a control and an experimental group is a standard approach to characterize the response of the circadian clock system to a treatment. Many such studies in high-impact journals discovered large-scale circadian changes resulting from the treatment and often described this phenomena as “circadian reprogramming” or “circadian remodeling”. VDA was used in most studies to determine the number and identity of DiffR features. VDA finds rhythmic transcripts in the two groups separately and then compares the lists of rhythmic transcripts to separate DiffR from non-DiffR transcripts. Analyses of high-throughput data stringently control for false discoveries at the cost of missing many true discoveries. In this work, we questioned the validity of the VDA due to the propagation of errors (false discoveries and missed true discoveries) from two separate rhythmicity tests to the determination of DiffR transcripts.

We showed using two artificial experiments constructed from real liver transcriptomic data that the VDA produces excessive false positive DiffR hits (Fig. 1 and Supplemental Fig. S1). VDA misidentifies large numbers of DiffR transcripts in datasets both with and without true DiffR transcripts. Irrespective of the stringency of rhythm detection, VDA manages to recover only a fraction of DiffR transcripts with a low accuracy in prediction as well. Even with the optimal choice of FDR threshold, VDA recovered only 30% of DiffR transcripts with 80% of the hits being incorrect. For the standard FDR threshold of 0.05, VDA recovered 40% of DiffR transcripts, but with only a 10% of hits being correct. The failings of VDA cannot therefore be sidestepped by being very conservative (very low FDR threshold) or being permissive (larger FDR threshold) with rhythm detection in either group. This issue with VDA in fact generalizes beyond transcriptomes to any high-throughput data and any effect (not just rhythmicity) measured independently in the two datasets.

We presented a model selection and a hypothesis testing approach to identify DiffR features (Fig. 2). Both these approaches called utmost a handful of DiffR hits for the first scenario with no true DiffR transcripts. For the second scenario with known true DiffR transcripts, these approaches identified nearly perfectly between 50-75% of the DiffR transcripts depending on the choice of algorithmic parameters. We explored pre-filtering candidate DiffR transcripts with a minimum amplitude for rhythmic transcripts (Fig. 3). An amplitude threshold for rhythms reflects one’s definition of what constitutes *biologically relevant* rhythms (Lück and Westermark, 2016). With an amplitude threshold, both approaches recover a high fraction of such biologically-relevant DiffR transcripts (with amplitude in either group exceeding the chosen threshold) at a slight loss of precision. Both approaches performed well, but model selection had low precision (higher number of false hits) in some situations.

We provide both approaches and their implementations for different data in a convenient to use R package *compareRhythms*. The default settings in the package correspond to the best performance tradeoff between precision and recall (according to the second scenario). The approaches however differ in more than just performance. We summarize the trade-offs involved in Table 1. The choice of approach must consider the nature of the data (transcriptomic vs. non-transcriptomic data), covariates, effects of experimental batches, waveforms of interest (sinusoidal vs. non-sinusoidal), size of the datasets/speed and experimental design complexity.

**Table 1:** Advantages and disadvantages of the analysis pipelines in *compareRhythms.* Linear modeling encompasses all implementations of hypothesis testing other than *DODR*, i.e., *limma, voom, DESeq2, edgeR*.

| Hypothesis testing |  | Model selection |
| --- | --- | --- |
| *DODR* | linear modeling |  |
| + straightforward | + very fast | + simple, elegant and fast |
| + rhythmic detection using RAIN | + blends easily to standard pipelines for transcriptomic data | + directly provides all DiffR categories |
| + works on any normalized data | + can include other covariates and batch variables | + works on any normalized data |
| – mixes parametric (for DiffR detection) and non-parametric methods (for rhythm detection) | – rhythm and DiffR detection based on sinusoidal rhythm pattern | + can include other covariates and batch variables |
| – slow |  | – exponential growth in models with rhythm patterns of interest |
| – cannot include other covariates in pipeline |  | – slightly worse performance |

**Table 2:** Accession number of the public data from GEO analyzed in this study.

| Publication | Accession number | Data type |
| --- | --- | --- |
| Hughes et al. (2009) | GSE11923 | microarray |
| Eckel-Mahan et al. (2013) | GSE52333 | microarray |
| Quagliarini et al. (2019) | GSE108688 | RNA-seq |
| Tognini et al. (2017) | GSE87425 | microarray |
| Pei et al. (2019) | PRJNA449625 | RNA-seq |

Model selection and hypothesis testing classify DiffR transcripts into four categories: transcripts that gain rhythms, lose rhythms, have the same rhythms or whose rhythms have changed (amplitude and/or phase) between the control and experimental groups. VDA analysis completely disregards the last category of rhythms. The apparently intuitive set theoretic approach underlying VDA only demarcates three groups from two lists of rhythmic transcripts (sets). While Venn diagrams are not themselves the problem, they are often symptomatic of the use of the flawed VDA and are better avoided.

To evaluate our conclusions from artificial scenarios further, we reanalyzed three circadian studies that used VDA; we also included a fourth study that repeated one of the studies without reporting a DiffR analysis. These studies focused on the effect of metabolic changes caused by dietary or genotype alterations on the mouse liver circadian transcriptome. For comparison, we used the reported results of the flawed VDA analysis in these studies. Across all studies, hypothesis testing and model selection called between 5-23% and between 34-51% of rhythmic transcripts to be DiffR, respectively. The highest fraction of rhythmic transcripts were DiffR in response to KD, followed by disruption of the H_2_O_2_ rhythm and then with HFD, and the ordering was independent of the approach used. The two studies on the effect of HFD contained very similar fractions of DiffR transcripts despite differences in assay and lab. However, VDA in the original studies reported upwards of 72% of rhythmic transcripts were DiffR. Considering the very high precision of hypothesis testing and a conservative estimate of its recall of 50% (Fig. 3), the number of true DiffR transcripts in these studies is at most 2 times our reevaluated hypothesis testing-based numbers. Taken together, all these studies significantly overestimate the extent of reprogrammed circadian rhythms consistent with our evaluation of the VDA approach they employed.

The identity of DiffR transcripts is as important as the size of the DiffR transcriptome and it allows for functional interpretation. To that end, we quantified the discrepancy between the DiffR hits from hypothesis testing to those reported in the original studies. Between 43-73% of DiffR hits called by hypothesis testing were absent in the DiffR hits called by VDA. These misses included DiffR transcripts with altered circadian parameters in the ‘change’ category, which is absent in the VDA and several that were called arrhythmic by VDA. We used calls from hypothesis testing for this comparison, since model selection appeared to have lower precision based on the artificial scenario. In fact, DiffR hits from model selection were a superset of the DiffR hits from hypothesis testing across all studies. Model selection behaves like hypothesis testing with a more liberal FDR threshold. In summary, VDA not only overestimates the number of DiffR hits but also overlooks a significant fraction of relevant DiffR transcripts.

We next explored whether this large divergence in the size and identity of the DiffR transcripts between our analyses and VDA affected the conclusions drawn in the original studies. Under HFD, the small set of DiffR transcripts still supported the original conclusions that many transcripts lost but none gained rhythms, two-thirds of transcripts had reduced amplitudes and almost all, not just a majority, of DiffR transcripts were phase-advanced (Fig. 4B,E). Although a small DiffR set precluded meaningful enrichment analysis, we observed tendencies that ‘circadian rhythms’ and ‘metabolic pathways’ were important (Fig. 4D). Under KD, we could corroborate the gain of rhythms in a plurality of DiffR transcripts (Fig. 5A). Moreover, we found no KEGG enrichment of ‘metabolism’ or ‘PPAR signaling’ among DiffR transcripts in the liver (Fig. 5C). Finally, in opposition to the original study, DiffR transcripts (with changed rhythms) showed no trend in the liver (Fig. 5B). On altering H_2_O_2_ rhythms, more transcripts gained and changed rhythms than lost rhythms (Fig. 6A). The original study found equal numbers gained and lost rhythms. We could confirm that DiffR transcripts that changed rhythms had altered phase, but not that a majority of these were phase delayed; we found no phase preference. Finally, we found no enrichment of ‘oxidation-reduction process’ or ‘metabolic process’ among DiffR transcripts (Fig. 6C). To sum up, we found some conclusions held up under the reanalysis, some did not and we were unable to evaluate yet others. In other words, high-throughput circadian studies must be re-assessed individually.

We wondered next whether our reanalysis produced novel hypotheses overlooked in the original studies. We conclude based on both studies that the effect of HFD is restricted to a relatively compact set of transcripts including the entire core clock network. Further, DiffR transcripts show a very consistent ~4h phase advance of rhythms (Fig. 4E). We hypothesize that *Nampt* with the same phase advance (Fig. 4C) and slight amplitude decrease drives a similar pattern in the metabolome (The metabolome analysis using VDA can be easily redone using *compareRhythms*). We have strong evidence based on KEGG enrichment and hallmark gene set analysis that KD activates viral defense and Interferon *α, γ* response pathways by inducing *de novo* rhythms in genes involved in these pathways (Fig. 5D). A recent study (Goldberg et al., 2019) showed that KD provides protection against influenza infection and we hypothesize that this effect is at least partially mediated by the circadian system. Finally, reactive oxygen species play a complex role in cancer (Reczek and Chandel, 2017). We found significant overlap between DiffR transcripts and transcripts upregulated in response to KRAS signaling based on hallmark gene sets (Fig. 6C,D). We propose that this interaction between redox balance and cancer is mediated by the circadian clock. Our analysis also suggests a relationship between redox balance, circadian clock and blood vessel development. Our novel insights thus related to the modulation of the interaction between physiological processes by the circadian network.

Our conclusions nonetheless are limited by the assumptions and formulation of our approach. The changes in amplitude and/or phase used to find DiffR transcripts are estimated assuming sinusoidal rhythm patterns. This assumption might be unsuitable in specific situations (e.g., long/short photoperiods). We also used a minimum rhythm amplitude for inclusion of rhythmic features in our analysis, because we believe this extracts more biologically meaningful results. This might lead to smaller DiffR sets in certain datasets. Removing this prefiltering in fact *reduced* the fraction of rhythmic transcripts that were DiffR in all three studies (Figs. S3A, S4A, S6A). Prefiltering improves the power of the subsequent DiffR test. In two of three studies, the number of DiffR transcripts was also smaller. Interestingly, without an amplitude threshold, hypothesis testing can only classify transcripts as ‘change’ and ‘same’. An amplitude threshold is necessary to define arrhythmicity in one group. This raises the question whether the absolute number of DiffR transcripts or the fraction of DiffR transcripts among all rhythmic transcripts is consequential.

We only reevaluated transcriptomic datasets relating the circadian clock and metabolism in the mouse liver due to their sheer abundance in public archives. However, our tool can be easily applied to any normalized (according to the particular datatype) data, such as metabolomic and proteomic data, from any organism. We recommend on the basis of this work that studies use our tool *compareRhythms* or a similar conceptual pipeline to perform DiffR analysis and to reevaluate old studies on a case by case basis. This tool is currently designed to only compare two groups with one categorical variable to account for covariates, such as batch or sex. In addition, we also disregard changes in mean expression between the two groups. Such more complex experimental designs (for e.g., Ananthasubramaniam et al., 2018) can be handled using a customized pipeline using other tools, such as *LimoRhyde* (Singer and Hughey, 2019) or *dryR* (https://github.com/naef-lab/dryR). We lent towards simplicity and performance in *compareRhythms* at the cost of including many different experimental designs.

The reproducibility crisis in science is widely accepted and problems with statistics and experimental design is often cited as one of the main culprits (Baker, 2016). The deficiency of the common approach to DiffR analysis is related to a common mistake of comparing two experimental effects without directly comparing them (Nieuwenhuis et al., 2011; Makin and Orban de Xivry, 2019). Venn diagrams that are symptomatic of VDA ought to serve as a warning flag. We trust that chronobiologists will find our tool an easy way to avoid this pitfall and generate reliable hypotheses to best utilize their resources.

## Methods

All analysis and statistics were performed using R 3.6.3 (R Core Team, 2020).

### Data sources

All data used in this study were gathered from Gene Expression Omnibus (GEO) database (Barrett et al., 2012) or the Short Read Archive. The accession numbers for the different studies are:

### Data Preprocessing

Raw microarray data were loaded using custom chip definition file from Brainarray (v24.0.0, Dai et al., 2005) with probes arranged and annotated according to Ensembl gene ID. They were subsequently normalized using the RMA algorithm in the *oligo* package (v1.50.0, Carvalho and Irizarry, 2010) to obtain final log_2_ expression values. The included subset of gene IDs had a minimum log_2_ expression of 4 in at least 70% of the samples in each condition.

Raw RNA-seq reads were quantified using *salmon* (v1.1.0, Patro et al., 2017) with *Mus musculus* reference genome (GENCODE build M24, Frankish et al., 2019). The salmon-quantified transcript expression was converted to gene expression using *tximport* package (v1.14.2, Soneson et al., 2015). We retained for the analysis all genes that had at least 10 mapped reads per 1 million reads in at least 70% of samples in each condition.

### Artificial scenarios

The 500 DiffR transcripts for the second scenario were created as follows: we selected 500 transcripts with harmonic regression (sine fits) adjusted p-values below 0.05 and peak-to-trough amplitudes above 0.26 (the median amplitude of rhythmic transcripts). We altered the samples in the odd group for each of these transcripts by scaling their mean-subtracted expression values by a uniform random variable in [0,1]. We then circularly shift the samples in the odd group for each transcript a number of places chosen randomly in [1,24]; there are 24 time-points in the odd group. Finally, we added Gaussian noise with standard deviation of ¼ the transcript amplitude to the odd group of samples.

### DiffR Identification

The hypothesis testing and model selection approaches are outlined in Fig. 2 and in the text and were implemented in the package *compareRhythms* (https://github.com/bharathananth/compareRhythms/). Expression values of the expressed genes were processed with default parameter values using *limma* (Ritchie et al., 2015), *DODR* (Thaben and Westermark, 2016) or model selection (Ritchie et al., 2015) pipelines for microarray and voom (Liu et al., 2015) or DESeq2 (Love et al., 2014) pipelines for RNA-seq data. For the hypothesis testing-based approaches, all p-values were false discovery adjusted using Benjamini-Hochberg correction and adjusted p-values were thresholded at 0.05 (unless otherwise mentioned). The amplitude threshold was set at 0.5 log_2_ expression with amplitude measured peak-to-trough.

The implementation of VDA for the two artificial scenarios was based on rhythm detection using RAIN (Thaben and Westermark, 2014) with the resulting p-values adjusted using the Benjamini-Hochberg approach. For VDA with amplitude threshold, the peak-to-trough amplitudes were determined using harmonic regression of the normalized expression values.

### Gene Enrichment

Gene enrichment analysis was performed using the *clusterProfiler* package (v3.14.3, Yu et al., 2012) with KEGG database (Kanehisa et al., 2017) and *msigdbr* (v7.1.1, Dolgalev, 2020) package translation of Molecular Signatures database (Liberzon et al., 2015).

## Code Availability

The full reproducible code as an R-markdown file to generate all the figures is available in the Supplemental Materials.

## Acknowledgments

This work was supported by Deutsche Forschungsgemeinschaft (DFG, German Research Foundation) grant AN 1553/2-1 to Bharath Ananthasubramaniam and DFG - Project-ID 278001972 - TRR 186 to Hanspeter Herzel and Achim Kramer. The authors thank Marta del Olmo for useful comments on the manuscript.

## Author contributions

B. A. designed the study. B. A. and A. P. performed the analyses and analyzed the results. B. A. wrote the manuscript. B. A., A. P., A. K. and H. H. edited and approved the manuscript.

## Disclosure Declaration

The authors declare no conflict of interest and no competing financial interests.

## Notes

### Competing Interest Statement

The authors have declared no competing interest.

